# SUMO – *In Silico* Sequence Assessment Using Multiple Optimization Parameters

**DOI:** 10.1101/2022.11.19.517175

**Authors:** Andreas Evers, Shipra Malhotra, Wolf-Guido Bolick, Ahmad Najafian, Maria Borisovska, Shira Warszawski, Yves Fomekong Nanfack, Daniel Kuhn, Friedrich Rippmann, Alejandro Crespo, Vanita Sood

## Abstract

To select the most promising screening hits from antibody and VHH display campaigns for subsequent in-depth profiling and optimization, it is highly desirable to assess and select sequences on properties beyond only their binding signals from the sorting process. In addition, developability risk criteria, sequence diversity and the anticipated complexity for sequence optimization are relevant attributes for hit selection and optimization. Here, we describe an approach for the *in silico* developability assessment of antibody and VHH sequences. This method not only allows for ranking and filtering multiple sequences with regard to their predicted developability properties and diversity, but also visualizes relevant sequence and structural features of potentially problematic regions and thereby provides rationales and starting points for multi-parameter sequence optimization.

## 1 Introduction

Due to the nature and setup of antibody and VHH display campaigns, identified screening hits generally show high antigen-specific binding. Usually, such sequences need to be optimized to fulfill all required criteria with respect to the targeted developability profile. Typical aspects that need to be considered to convert an antigen-specific binder into a developable molecule include the assessment and reduction of the following risk factors: immunogenicity, chemical liabilities and post-translational modifications (PTMs), physical instabilities, viscosity, poly-specificity or poor expression. These aspects may not only affect the pharmacodynamic and pharmacokinetic profile of the drug substance, but also hinder the manufacturing and development process of the drug product. Sometimes, the complexity of these different developability parameters might require multiple design cycles and in some cases it might not be even possible to optimize such hits towards a favorable overall profile [1]. Therefore, it would be highly desirable to select sequences obtained from display campaigns not only on the basis of their binding signals from the sorting process, but also based on their overall developability profiles and anticipated complexity for multi-parameter optimization.

In recent years, progress has been made to implement high-throughput predictive experimental developability assays with low compound need and short cycle times that can be used for efficient early profiling and identification of potential developability risks [2–8]. However, these *in vitro* methods still require resources and are therefore not always suited to filter and rank large sequence sets that might physically not be available in sufficient amounts in the early hit finding and optimization phase.

In analogy to Lipinski’s rule of five to prioritize the selection of small molecules for entry into clinical development [9], several metrices have been implemented for an *in silico* developability assessment of antibody sequences [10, 11]. One of the first structure-based approaches is the “spatial aggregation propensity” (SAP) [12] score, that was later combined with the antibodies net charge into the Developability Index [13]. Another approach, implemented as solubility predictor, is CamSol [14], which was also used to design antibody variants towards an improved developability [15]. In recent years, further approaches that investigated the *in silico* property distribution of clinical or marketed antibodies to identify the most descriptive and informative properties have allowed to assess whether a sequence might be developable or not [16–18]. In those studies, three-dimensional (3D) models of the antibodies were generated and used as input for the calculation of different structure- or sequence-based descriptors. In each study, five different descriptors were identified that include combinations of hydrophobic and electrostatic descriptors. In detail, the Therapeutic Antibody Profiler (TAP) [18] provides CDR length, surface hydrophobicity in CDR vicinity, positive and negative charge patches in CDR vicinity and a structural Fv charge symmetry parameter as key *in silico* descriptors. An in-depth study of the intrinsic physicochemical profile of marketed antibodies from Boehringer Ingelheim [16] yielded the following nonredundant descriptors as potential developability criteria: surface area buried between the variable light (VL) and heavy (VH) domains, the isoelectric point (pI), ratio of dipole moment to hydrophobic moment, ratio of surface areas of charged to hydrophobic patches and a scale for hydrophobic imbalance. Recently, the Therapeutic Antibody Developability Analysis (TA-DA) [17] approach suggests the following scores to efficiently separate clinical antibodies from repertoire antibodies: AggScore [19] predictions of the light chain framework1 region, AggScore predictions of the framework region of the entire antibody, positive patches of the light chain CDRs, an atomic contact energy that reflects burial of hydrophobic residues and exposure of charged and polar atoms and TOP-IDP, a summed amino acid scale reflecting the propensity for intrinsic disorder.

These *in silico* scores reflect general physical stability attributes of antibody sequences and seem well suited for filtering and ranking sequences in terms of their manufacturability and solution behavior, e.g. viscosity and colloidal stability. It was also found that the same hydrophobicity and electrostatic property descriptors are useful for identifying sequences, which might lead to fast *in vivo* clearance via different mechanisms as outlined below.

In addition to these physical stability aspects, chemical (e.g. deamidation, isomerization or oxidation) or post-translational modifications (PTMs, such as glycosylation) represent a further risk factor for developability [7, 20]. Such modifications could be introduced in the manufacturing process, during storage in the administration device or after *in vivo* administration and can result in species with reduced binding affinity or might even trigger an immunogenic response. Several *in silico* approaches have meanwhile been reported that allow to predict such chemically labile or post-translational modification sites, either based on the sequence or considering structural properties in addition [21–26].

Furthermore, it is well known that immunogenicity can also be triggered by the native antibody sequence itself. Non-human antibodies have been demonstrated to induce human immune responses, often resulting in neutralization of the administered antibody and causing enhanced clearance. Humanized and even fully human sequence-derived antibody molecules can still carry an immunological risk [27]. A common approach to reduce this risk is the framework-based germline humanization of antibody sequences [28]. Moreover, a common practice for an early immunogenicity assessment of biotherapeutics is the *in silico* prediction, identification and de-immunization of potential non-self regions of the antibody sequence that can bind to the major histocompatibility complex (MHC) on antigen presenting cells. The main output from such *in silico* based risk assessment includes the predicted binding affinity of epitopes binding to MHC class II alleles and the promiscuity of MHC class II alleles that have high affinity epitopes predicted to bind [29, 30].

With these *in silico* approaches available in the public domain or in commercial software vendors, we aimed to implement a workflow that allows to provide a lean *in silico* assessment regarding the above-mentioned developability aspects for antibody or VHH sequences. This workflow, called **SUMO** (*In Silico* **S**equence Assessment **U**sing **M**ultiple **O**ptimization Parameters), automatically computes 3D models and uses different software packages to calculate a diverse set of more than 250 physico-chemical sequence- and structure-based *in silico* properties. From these, four key properties (pI; Positive Patch Energy of the CDRs; AggScore of the variable region; and AggScore of CDR regions) were selected as electrostatics and hydrophobicity descriptors, based on their orthogonality (as described by Ahmed et al. [16]), general interpretability and since they reveal good direct correlation to several relevant experimental developability parameters. The pI, for example, has been shown to be a useful filter criterion regarding colloidal stability in standard formulations [5, 31] and non-specific binding that might lead to fast clearance [16, 32]. In addition, it was demonstrated in several studies that highly positively charged CDRs can contribute to fast clearance by different mechanisms [6, 33–42]. AggScore [19] penalizes clusters of surface-exposed hydrophobic atoms that do not have surrounding charge patches to mitigate inter-molecular hydrophobic-hydrophobic interactions and has shown good predictivity to internal HIC (hydropbobic interaction chromatography) data. High hydrophobicity is known to be problematic for non-specific binding (potentially leading to fast clearance), expression yields and solution behavior (viscosity, colloidal stability) in high concentration formulations. In addition to the computed AggScore for the entire Fv region, **SUMO** also reports the AggScore derived from the CDR regions, to indicate whether removal of such hydrophobicity/aggregation spots requires amino acid modifications in the CDRs. In addition to these electrostatic- and hydrophobicity-related parameters, the overview **SUMO** report provides information on: (i) the number of predicted chemical liabilities (deamidation, isomerization and methionine oxidation sites) and PTMs (N-linked glycosylation sites) within the CDRs; (ii) sequence identities of the framework region and full-length variable regions compared to the most similar human germline sequences; and (iii) an *in silico* immunogenicity assessment indicating whether or not the strongest and most promiscuous 15-mer peptide of the antibody/VHH sequence is present in the human germline repertoire. Finally, for larger sets of sequences, clustering information is also provided (for different regions of the antibody), to assess sequence diversity. These *in silico* scores are summarized in an overview table (with one sequence per row) that includes a color coding (green-yellow-red) to support straightforward visual assessment of the sequences based on their computed *in silico* developability profiles.

Whereas this overview table (Figure 1) allows to compare and filter different sequences with regard to their computed developability profiles, **SUMO** also generates sequence views for each antibody or VHH with a detailed assessment for each residue along the amino acid sequence (Figure 2). This detailed assessment includes different antibody annotation schemes (IMGT, AHo, CCG, Chothia and Kabat), a sequence alignment to the most similar germline sequences, the information whether a residue is located in a key structural position (e.g., Vernier, VL/VH Interface), whether the residue is a sequence- or structure-based predicted liability or glycosylation site and the computed per-residue surface exposure, surface hydrophobicity, charge, solubility and aggregation scores (vide supra). Furthermore, MHC-II binding scores of all 15-mer peptides along the amino acid sequence against a reference panel of 27 HLA alleles is provided. The scores are complemented with a color coding that allows to visually identify hot spot regions for each specific optimization parameter.

**Figure 1.**
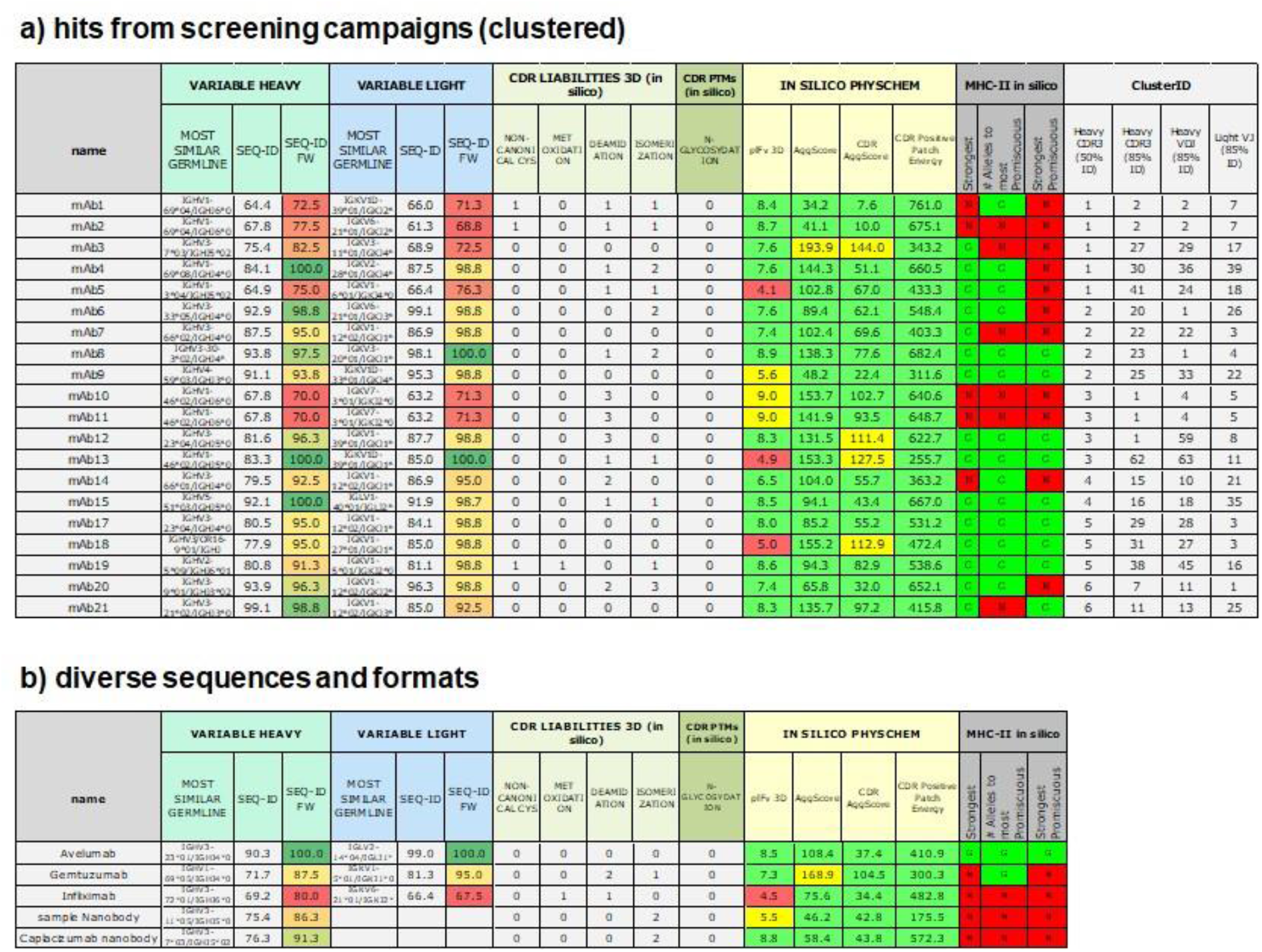
**SUMO** overview table over different antibody/VHH sequences. a) Multiple antibody sequences from a screening campaign with the following *in silico* properties reported for each sequence: (i) name of the most similar germline sequence and the sequence identify based on the entire variable chain or the framework (FW) region only, (ii) the total number of specific chemical liabilities and PTMs (non-canonical cysteines, methionine oxidation, deamidation or isomerization, N-glycosylation) in structurally exposed CDR residues, (iii) the structure-based pI and AggScore of the variable region, AggScore of CDR regions only and the Positive Patch Energy of the CDRs as physico-chemical developability descriptors and (iv) predicted MHC-II binding flags that indicate whether the strongest and most promiscuous 15-mer peptide epitopes occur in the human germline repertoire: G (Germline, green); N (Non-germline, red). The sequences are sorted based on their cluster IDs, thereby facilitating the selection of sequences based on computed *in silico* properties and diversity. These computed physico-chemical properties (iii) are complemented with a color coding as outlined in **Note 3**. b) Additional examples of marketed antibodies and VHHs (Nanobodies) to illustrate the applicability of SUMO regarding sequence and format diversity.

**Figure 2.**
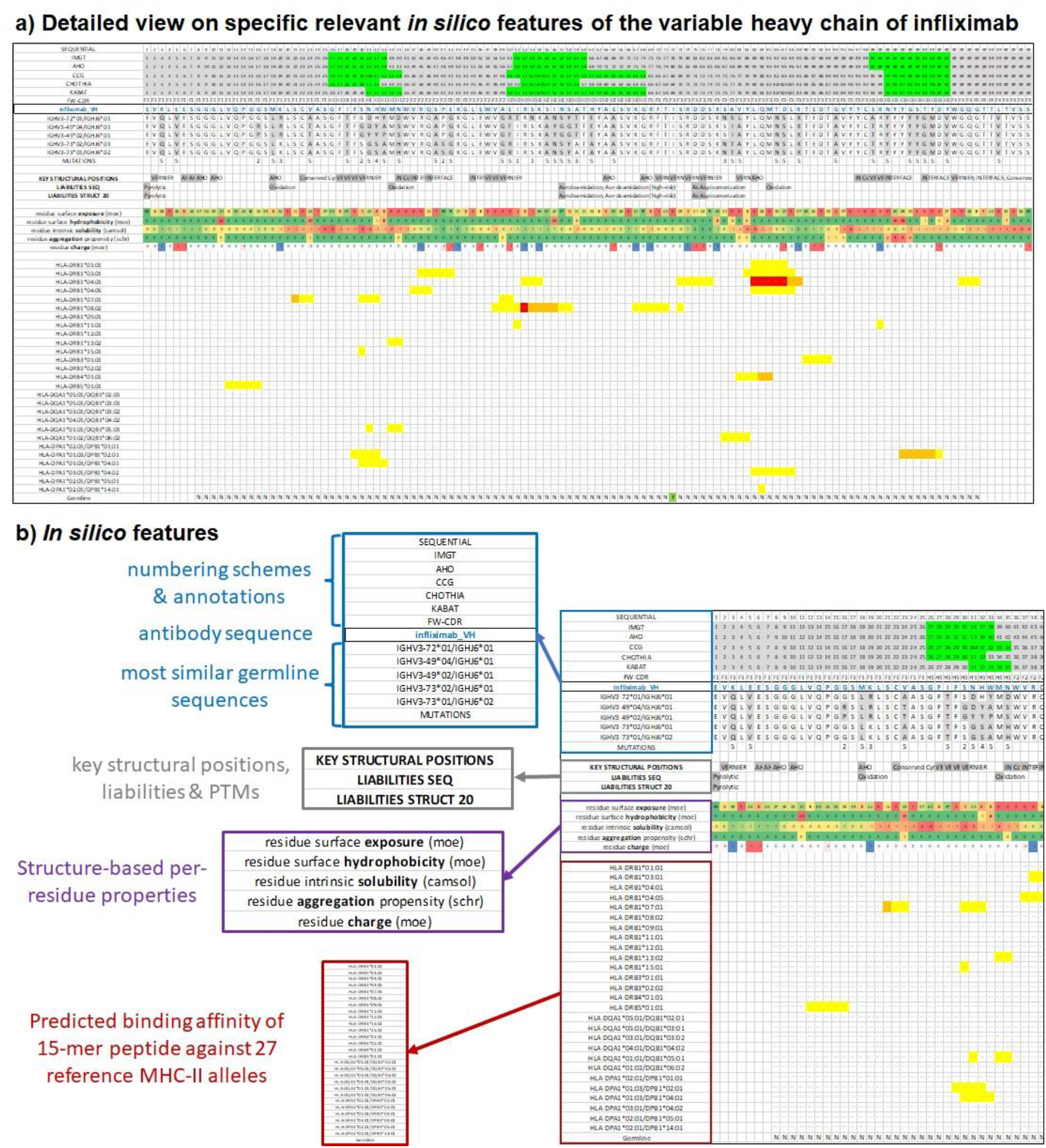
Detailed **SUMO** tables for variable light and heavy chains are generated for each sequence. a) Detailed view visualizing specific relevant features of the variable heavy chain sequence of Infliximab as an example. Each column is devoted to a residue in a sequence, which are aligned to the most similar human germline sequences, annotated and complemented with specific per-residue developability scores. b) Schematic explanation of *in silico* features (shown in rows) for each residue of the infliximab variable heavy chain: (i) Sequence numbers according to different numbering schemes. The positions of CDRs are highlighted in green. (ii) Residues of the most similar germline sequences with amino acid differences shown in gray. (iii) Information about “key structural positions” and sequence-as well as structure-based predicted liabilities and PTMs. (iv) Structure-derived per-residue surface exposure, surface hydrophobicity, CamSol predicted solubility, Aggregation Scores (AggScore) and charge, including a color coding. (v) predicted binding strength for each 15-mer peptide along the amino acid sequence against a reference set of 27 MHC-II alleles. The central amino acid of the 15mer is highlighted in red, orange and yellow according to the predicted binding strength.

Finally, **SUMO** also provides 3D visualizations of the computed properties on the automatically generated antibody/VHH models via PyMOL session files that can be accessed via PyMOL scenes for easy structural interpretation (Figure 3).

**Figure 3.**
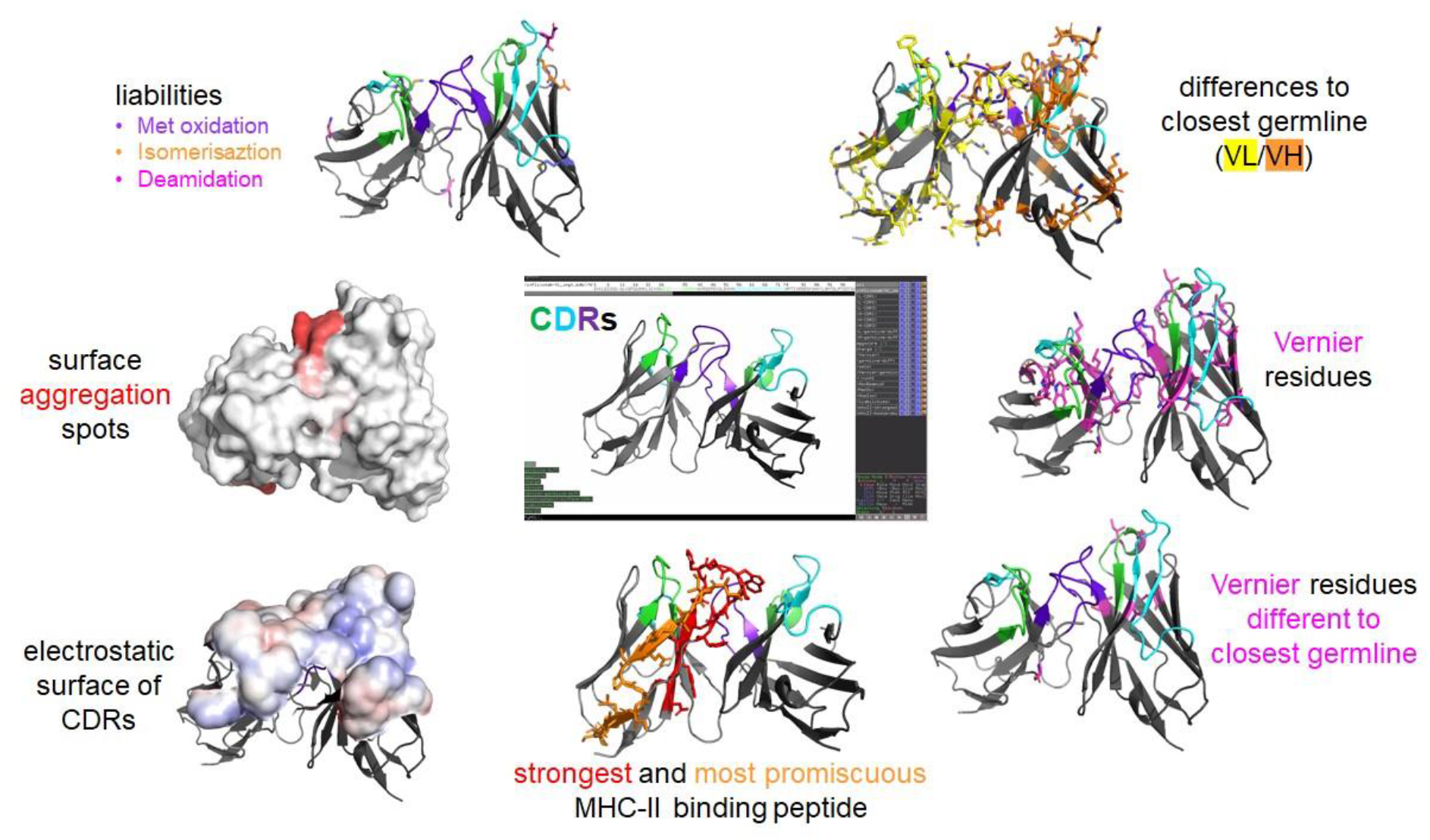
Illustration of different scenes in PyMOL session files automatically generated from SUMO tables and homology models. Visualization of specific molecular properties and potential risk factors that are relevant for sequence assessment and optimization in 3D is a powerful tool to visually analyze and assess the potential areas of concern. The scenes can be displayed interactively, views can be rotated, high resolution graphic views can be exported and shared with colleagues.

The presented **SUMO** workflow has been automated and requires as input (only) a list of FASTA sequences of the variable region of antibodies or VHHs. This approach can provide a very early *in silico* developability risk assessment already at the sequence level for several parameters that can experimentally only be determined late in drug discovery programs, such as physical stability attributes, immunogenicity or pharmacokinetic properties. This early *in silico* assessment can be used to (i) support the selection of hit sequences from display campaigns in addition to the available experimental binding data, (ii) trigger and prioritize experimental testing of sequences in predictive *in vitro* developability assays based on the identified *in silico* risks and (iii) to design sequences towards a favorable overall developability profile. Taken together, this early *in silico* sequence assessment has the potential to accelerate candidate selection and reduce risks and attrition rates in clinical development.

## 2 Materials

1. FASTA files containing variable domain sequences of antibodies were provided as input.
2. Molecular Operating Environment (MOE) [43] 2020.0901 by Chemical Computing Group was used to build antibody and VHH models, *in silico* property calculation and antibody annotation.
3. IgBLAST [44] was utilized to identify and align the most similar human germline sequences.
4. Schrodinger/BioLuminate 4.6 [45] was used for antibody/VHH *in silico* property calculations.
5. CamSol (version 2.1) [14] was used to predict the intrinsic solubility profile of sequences.
6. An inhouse implementation of the Therapeutic Antibody Profiler scores (TAP-Scores, Raybould et al., 2019).
7. The Immune Epitope Database and Analysis Resource (IEDB) software (version IEDB-3.1.6) [46] was used to predict MHC-II binding of antibody/VHH peptide fragments.
8. IMGT/V-QUEST reference set was used to identify human versus non-human germline sequences [47–49].
9. Geneious Biologics [50] was used for sequence clustering of antibody/VHH sequences.
10. PyMOL [51] was used for structural visualization.
11. Several inhouse scripts were provided for analysis and post-processing of the workflows.

## 3 Methods

### 3.1 Sequence annotation and calculation of in silico properties

The first step includes the preparation of the variable domains of antibody or VHH sequences, the annotation according to different numbering scenes, the identification of the most similar human germline sequences, the generation of 3D models and the calculation of sequence- or structure-based global and per-residue descriptors using different software packages.

1. Provide single FASTA files containing light and heavy variable region chains of antibodies or heavy variable region chains of VHHs. With these as input, the following steps are executed automatically by a pipeline of Python (version 3.8) and Scientific Vector Language (SVL) [43] scripts.
2. FASTA file is split to create individual FASTA files per antibody
3. FASTA files are imported into MOE along with user defined ForceField configuration to generate homology models using the Antibody Modeler. MOE project database is regularly updated to include internal antibody crystal structures data. To optimize the analysis time, the number of models is set to 1, when analyzing screening campaigns (*see* **Note 1**).
4. Assignment of numbering schemes for each sequence (IMGT, AHo, CCG, Chothia and Kabat).
5. Identification of most similar human germline sequences and calculation of sequence identities over the entire variable region and the framework regions based on the selected annotation scheme (by default: IMGT).
6. Identification of sequence- and structure-based (default solvent exposure cutoff: 20%) liabilities or optionally sequence-only liabilities such as unpaired cysteines, methionine oxidation, deamidation, isomerization, N-linked glycosylation sites by MOE.
7. Global and per-residue *in silico* property (descriptor) calculations. Three-dimensional (3D) homology models of the antibody or VHH variable domains are provided as input to calculate a total of 251 global structure- and sequence-based descriptors within (i) MOE including BioMOE, (ii) Schrodinger/BioLuminate, (iii) CamSol or (iv) TAPscore.

### 3.2 MHC-II Binding Predictions

For an *in silico* immunogenicity assessment, the antibody or VHH sequence is automatically split into all possible 15-mer peptides along the amino acid sequence. For each peptide, (i) a binding score to a reference set of MHC class II alleles is computed (using the IEDB software) and (ii) its occurrence in the human germline repertoire is investigated.

1. Provide the full amino acid sequences (including the constant regions) as FASTA file as input and run the IEDB MHC-II binding predictions using the following settings as input: IEDB recommended Prediction Method; Selected species/locus: Human, HLA-DR; the IEDB-recommended reference panel of 27 alleles; peptide length: 15.
2. Consider all 15-mer peptides among the top 5% in percentile rank as binders, among the top 2% as strong binders, and among the top 1% as very strong binders to MHC-II alleles.
3. Identify for each 15-mer peptide whether it occurs in the human germline repertoire (labelled as “G”) or not (labelled as “N”) by comparing to a copy of IMGT/V-QUEST reference set.
4. Generate an overview table that contains the predicted binding affinity of each 15-mer peptide against all considered MHC-II alleles.
5. Identify the predicted strongest and most promiscuous 15-mer peptides.
6. Peptides that are identified as strongest or most promiscuous MHC-II binders and do not occur in the human germline repertoire are considered as potential immunogenicity risk that might be subjected to sequence optimization towards de-immunization (*see* **Note 2**).

### 3.3 Sequence Clustering

To select sequences for hit evaluation not only based on their binding affinity and *in silico* developability profile, a list of sequences is subjected to sequence clustering. Considering sequence diversity as additional selection criterion increases the likelihood to identify hits that might bind to different epitopes.

1. Provide sequences of the variable regions to Geneious Biologics.
2. Perform sequence-identity based clustering using the following regions and thresholds: (i) H-CDR3, 50%, (ii) H-CDR3, 85%, (iii) H-VDJ, 85% and (iv) L-VJ, 85% and extract the cluster ID for each region.

### 3.4 Generation of an overview table on the in silico developability assessment of multiple antibody/VHH sequences

For each sequence, the following information is provided. This information serves as first assessment criteria regarding the general developability for an antibody/VHH sequence. For more detailed information on specific developability criteria, a detailed view for each sequence is provided (Method 3.5).

1. Report of (i) the most similar human germline sequence and sequence identity (ii) over the entire variable region and (iii) the framework regions only based on the selected annotation scheme (by default: IMGT).
2. Number of sequence- and structure-based (default solvent exposure cutoff: 20%) liabilities within the CDR regions.
3. From the set of 251 global structure- and sequence-based in silico descriptors, the following are reported: pI (of the variable domain), AggScore, CDR-Aggscore and CDR Positive Patch Energy. These scores are complemented with a green to yellow to red color coding (*see* **Note 3**).
4. From the computed MHC-II binding profile, provide a flag whether the strongest and most promiscuous binding peptide occurs in the human germline repertoire (G) or not (N) (*see* **Note 2**).
5. Add the cluster IDs based on the computed clustering regions and cutoffs. The table is sorted by these cluster IDs, thereby providing a view that has similar sequence clusters among each other.
6. A color coding is added (red to yellow to green) to the sequence identities vs the closest germline sequences that allows to assess and differentiate different sequences with regard to their human-likeness.
7. Optionally, any relevant experimental data is added to this table to allow for a ranking and prioritization of sequences based on the overall *in vitro* and *in silico* profile.

### 3.5 Generation of a detailed view on specific relevant features of the variable antibody sequence

In addition to this *overview table*, the SUMO workflow generates a detailed table for each specific sequence (see Figure 2). Inspection of this table is used for a specific assessment and evaluation of potential sequence liabilities and risks and serves as starting point for the design of variants towards sequence optimization, i.e., for humanization, de-immunization or elimination of liabilities or aggregation hot spots.

1. Display of the amino acid sequence and residue numbering using different schemes (IMGT, AHo, CCG, Chothia and Kabat). Frameworks and CDRs are annotated and highlighted (CDRs in green, Fig.2).
2. Display of sequence-aligned five most similar human germline sequences. Mutations, insertions/deletions are highlighted and summarized.
3. Information at the corresponding residue position, if this residue is located at a “key critical position” (Vernier, AHo, Interface, conserved cysteines).
4. Display at the specific positions if a residue is a sequence- and structure-based (default solvent exposure cutoff: 20%) predicted potential liability or post-translational modification site (i.e., non-canonical cysteine, methionine oxidation, deamidation, isomerization or N-linked glycosylation).
5. From the automatically generated antibody model (or x-ray structure, if available), the calculated per-residue (i) surface exposure, (ii) surface hydrophobicity, (iii) charge, (iv) CamSol predicted solubility and (v) Aggregation Score (AggScore) values are reported. To facilitate the identification of hot spots, these values are complemented with a red to yellow to green color coding.
6. MHC-II binding predictions of all 15-mer peptides along the amino acid sequence against a reference panel of 27 HLA alleles. The predicted binding scores of each 15-mer peptide (represented at the position of the central amino acid) are color coded (yellow to orange to red) by increased predicted binding strength. This visualization allows to identify all (non-germline) binders that are predicted to bind strongly to one or more MHC-II alleles and might be candidates for de-immunization.

### 3.6 Generation of PyMOL session files

Finally, based on x-ray structures or automatically computed three-dimensional (3D) models, the SUMO workflow automatically generates visualizations of relevant molecular *in silico* properties that are considered for a final structural assessment of relevant risk features and for the overall potential for multi-parameter “optimizability” (see Figure 3). Specific visualizations are stored as scenes in PyMOL session files and can be restored for visual inspection. For this purpose, a PyMOL script is automatically generated that performs the following steps:

1. Loads the pdb file of the antibody/VHH model or x-ray structure
2. Creates selections and specific-color codings for (i) CDR regions, (ii) predicted liabilities and PTMs, (iii) residues that are different from the closest germline sequence, (iv) key structural positions and (v) predicted strongest and most promiscuous MHC-II binding regions.
3. Creates color maps based on the values of the per-residue *in silico* properties: AggScore, Camsol, charge, residue-surface-hydrophobicity
4. Creates an Adaptive Poisson-Boltzmann Solver (APBS [52]) electrostatic surface map and a corresponding visualization for the CDR regions.
5. Saves different visualizations as individual scenes in a PyMOL session file.

## 4 Notes

1. Antibody structures are currently modeled using MOE’s antibody modeler, generating by default one conformation as output. It has been recognized that several predicted physico-chemical properties of antibody or VHHs show better correlations to experimental data when considering conformational ensembles instead of static structures, e.g. ref [53]. Our overall workflow has been modularized and therefore principally allows to generate structural models with alternative software tools and the usage of conformational ensembles for descriptor calculations.
2. Instead of reporting only the strongest and most promiscuous non-germline MHC-II binders, it is possible to provide alternative scores to assess the overall *in silico* immunogenicity risk, for example the number of (non-germline) MHC-II binders that are predicted to bind beyond a defined threshold score against any MHC-II allele. For a detailed assessment of the predicted MHC-II binding profile of specific antibody/VHH sequences, it is recommended to inspect the detailed overview table as described in Method 3.5.
3. The identification of cutoff scores for the *in silico* descriptors was based on their average values and standard deviations over a list of 79 Fvs from 77 biotherapeutics approved for human use, following the same procedure as described in ref (Ahmed et al., 2021). For the predictions of AggScore, CDR AggScore and CDR Positive Patch Energy, scores within one standard deviation from the mean are colored green, scores above one standard deviation yellow and those above two standard deviations red. For the AggScore scores, these cutoffs were slightly adjusted based correlation analyses to internal experimental HIC data. For the computed pI, the same color coding was applied to upper and lower deviations from the mean.

## 6 Acknowledgements

The presented approach is the result of a truly collaborative effort including many colleagues from different disciplines of our R&D organization. We would like to thank all these colleagues for continuous and constructive discussions, and (experimental and *in silico*) data acquisition that help to improve the predictivity of *in silico* predictions, and finally to establish **SUMO** as our standard workflow for early *in silico* sequence assessment for Biologics.

